# GPNMB is a biomarker for lysosomal dysfunction and is secreted via LRRK2-modulated lysosomal exocytosis

**DOI:** 10.1101/2025.01.01.630988

**Authors:** Erin C. Bogacki, George Longmore, Patrick A. Lewis, Susanne Herbst

**Author notes:** Laboratory of Neurogenetics, National Institute on Aging, Bethesda, MD 20892, USA. Corresponding author: Susanne Herbst.

## Abstract

Genome-wide association studies have identified *Glycoprotein Nmb* (*GPNMB*) as a risk factor for Parkinson’s Disease. The risk allele increases *GPNMB* transcription and GPNMB protein levels in the CSF highlighting GPMNB as a potential biomarker for Parkinson’s Disease. However, a lack of knowledge of GPNMB’s function and mechanism of secretion have hindered an interpretation of secreted GPNMB levels. In this study, we assessed the mechanism of GPNMB secretion by macrophages, the primary cell type expressing GPNMB in the brain. We show that GPNMB is secreted in response to lysosomal stress via lysosomal exocytosis and highlight the Parkinson’s Disease risk factor LRRK2 as a strong modulator of GPNMB secretion.

## Introduction

In recent years, our understanding of the genetic basis of Parkinson’s Disease (PD) has expanded with genome-wide association studies now highlighting over 90 gene loci which contribute to the development of the disease (Nalls et al. 2019). This expansion and refinement of the genetic architecture for PD has resulted in the new challenge of translating the genetic findings into therapeutic targets and biomarkers for this disorder.

One of the genes identified by genome-wide association is *Glycoprotein Nmb* (*GPNMB*), with the risk allele resulting in a 10% increased lifetime risk of developing PD. GPNMB, the protein product of *GPNMB*, is a transmembrane protein with a secreted extracellular domain. The GPNMB risk polymorphism results in increased GPNMB expression (Murthy et al. 2017; Diaz-Ortiz et al. 2022), and increased levels of the GPNMB secreted extracellular domain in the cerebrospinal fluid (CSF) (Brendza et al. 2021) – suggesting a role for increased secreted GPNMB levels in the pathogenesis of PD. In contrast, *GPNMB* loss-of-function mutations result in Autosomal-Recessive Amyloidosis Cutis Dyschromica (ACD), which manifests in abnormal skin pigmentation (Yang et al. 2018; Qin et al. 2021). GPNMB is homologous to melanosome amylogenic protein PMEL, and therefore is thought to play a role in melanosome biogenesis, which might underlie its role in the etiology of ACD (Van Der Lienden et al. 2018). In the context of PD, recent studies suggest that GPNMB can act as a receptor for alpha-synuclein uptake in neurons (Diaz-Ortiz et al. 2022). However, GPNMB KO failed to prevent the spread of alpha-synuclein or loss of dopaminergic neurons in several murine PD models (Brendza et al. 2021). Therefore, the pathogenic mechanism of either intracellular or secreted GPNMB in PD remains elusive. Intriguingly, GPNMB is highly expressed in microglia and has been described as a marker for pro-inflammatory microglia in PD (Brendza et al. 2021; Smajić et al. 2022).

GPNMB is under the transcriptional control of TFEB, a lysosomal stress-responsive transcription factor (Carey et al. 2020). This has led to GPNMB being proposed as a lysosomal protein aligning with a wider array of PD risk genes that highlight lysosomal dysfunction as a causal factor in PD (Bhore et al. 2024). For example, variation in the lysosomal enzyme GBA1 underlies common PD risk in diverse populations (Rizig et al. 2023; Nalls et al. 2019) and mutations in Leucine-rich repeat kinase 2 (LRRK2), which is recruited to stressed lysosomes and has been linked to lysosomal trafficking (Herbst et al. 2020; Bonet-Ponce et al. 2020b; Eguchi et al. 2018), represent one of the most common inherited form of PD. However, little is known about the physiological events that induce GPNMB secretion, or whether GPNMB function intersects with other lysosomal contributors to PD. These gaps in our knowledge hinder the interpretation of GPNMB as a secreted biomarker or the evaluation of GPNMB as a therapeutic target in PD.

This study investigated pathways modulating GPNMB secretion, revealing that dysfunctional lysosomes recruit GPNMB leading to the secretion of the GPNMB extracellular domain by lysosomal exocytosis. Linking *GPNMB* to monogenic forms of Parkinson’s, CSF levels of secreted GPNMB are increased strongly in *LRRK2* mutation carriers as well as *GBA1* mutation carriers and in idiopathic PD. These data suggest that secreted GPNMB can be considered a biomarker of lysosomal stress, and that this can reflect lysosomal dysfunction in genetic cases of Parkinson’s.

## Results

### Macrophages secret GPNMB in response to lysosomal stress

The *GPNMB* PD risk allele results in increased GPNMB CSF concentrations (Brendza et al. 2021) and previous reports have highlighted increased GPNMB levels in the plasma of PD patients (Diaz-Ortiz et al. 2022). To identify the source and potential triggers of GPNMB secretion in PD, we re-analysed a midbrain snRNA sequencing dataset of age-matched controls and idiopathic PD (iPD) patients (Smajić et al. 2022). *GPNMB* expression was the highest in microglia, followed by astrocytes and oligodendrocyte precursors (OPCs). However, *GPNMB* expression was only increased in iPD samples in microglia, pointing to microglia as a PD-responsive source of GPNMB (**Fig. 1A**). As previously reported (Brendza et al. 2021; Dolan et al. 2023), microglia subclustering identified damage-responsive microglia (DAMs) as the main subtype expressing *GPNMB* (**Fig. 1B-C**). Pathway enrichment analysis of genes differentially expressed in the DAM cluster highlighted dysregulation of cholesterol metabolism, lysosomal activation and hypoxic stress as potential microglia stressors. Congruent with this, *GPNMB* transcription is regulated by the MiT/TFE family of transcription factors (Carey et al. 2020; Loftus et al. 2009) indicating a role for GPNMB in the lysosomal stress response. To test the effect of lysosomal stress on GPNMB secretion, we first confirmed the transcriptional upregulation of GPNMB in response to various lysosomal stressors, such as the lysosomal damaging agent LLOMe, the lysosomal V-ATPase inhibitor Bafilomycin A1 and the K+/H+ anti-porter nigericin in RAW264.7 mouse macrophages. Lysosomal damage and alkalinization resulted in strong up-regulation of *GPNMB* transcription whereas Nigericin had a more modest effect (**Fig. 1E**). Additionally, we observed minor up-regulation of *GPNMB* gene transcription in response to proteotoxic stress induced by inhibition of the proteasome with MG132, but no transcriptional up-regulation after treatment with the TLR4 ligand LPS (**Fig. 1E**). Next, we assessed GPNMB secretion in mouse bone marrow-derived macrophages and RAW264.7 cells. In all cases, direct lysosomal stress resulted in GPNMB secretion whereas the effect of indirect stressors such as MG-132 varied. In contrast, LPS treatment did not result in GPNMB secretion. Overall, lysosomal damage induced by LLOMe and lysosomal alkalinisation with the V-ATPase inhibitor Bafilomycin A1 resulted in robust and substantial secretion of GPNMB (**Fig. 1F**). We also assessed GPNMB sub-cellular localisation in response to the same lysosomal and cellular stressors. Under steady-state conditions, GPNMB was predominantly localized to the Golgi and trans-Golgi (**Fig. 1H**). However, lysosomal stress resulted in the recruitment of GPNMB to LAMP1-positive lysosomes, indicating that GPNMB secretion is linked to lysosomal recruitment (**Fig. 1G-H**).

**Fig. 1.**
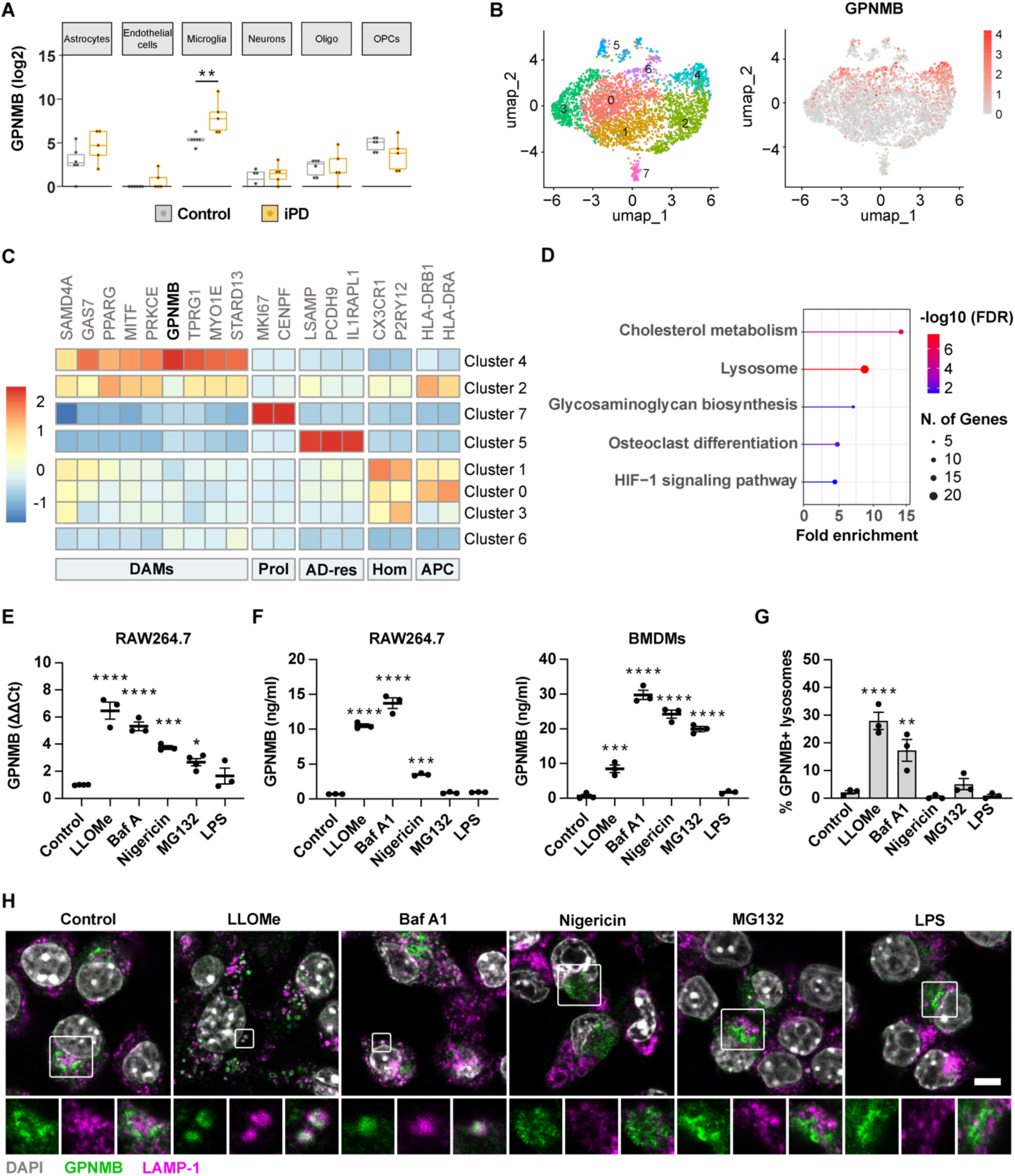
Macrophages secret GPNMB in response to lysosomal stress. **(A)** Average GPNMB expression across different cell types in the midbrain in controls and idiopathic PD cases. Analysed from (*6*). **(B)** Subclustering of microglia from (A) identifies a GPNMB high expression cluster.**(C)** Heatmap of average expression of genes associated with disease-associated microglia (DAMs), proliferating cells (Prol.), AD-resistant microglia (AD-res), homeostatic microglia (Hom.) and antigen-presenting microglia (APC). **(D)** Pathway enrichment analysis of DEGs of GPNMB-expressing DAM cluster. **(E)** RAW264.7 macrophages were either left untreated or treated with LLOMe (500 μM), Bafilomycin A1 (100 nM), Nigericin (5 μM), MG132 (5 uM) or LPS (10 ng/ml) for 4hrs and GPNMB transcription was assessed by qPCR. **(F)** RAW264.7 macrophages and BMDMs were either left untreated or treated with LLOMe (500 μM), Bafilomycin A1 (10 nM), Nigericin (1 μM), MG132 (5 uM) or LPS (10 ng/ml) overnight and GPNMB secretion was assessed by ELISA. (**G-H)** RAW264.7 macrophages were either left untreated or treated with LLOMe (1 mM for 1hr), Bafilomycin A1 (100 nM for 3 hrs), Nigericin (5 μM for 1 hr), MG132 (5 uM) or LPS (10 ng/ml for 1 hr) and GPNMB-LAMP-1 co-localisation was assessed by confocal microscopy. **(G)** Mean +/- SEM from three independent experiments. **(H)** Representative images, scale bar = 5 μm.

### GPNMB lysosomal recruitment is a pre-requisite for its secretion

To test whether GPNMB secretion is dependent upon lysosomal recruitment, we mutated motifs within the GPNMB cytosolic tail which are predicted to aid endolysosomal sorting. GPNMB S542 is the most robust phospho signal reported on Phospho Site Plus (Hornbeck et al. 2015), and forms part of a hypothetical export signal for sorting into clathrin-coated vesicles from the trans-Golgi (Owen, Collins, and Evans 2004). Y525 forms part of a hypothetical HemITAM motif required for AP-4 mediated clathrin-independent sorting and L562/L563 form the core of a Di-leucine motif, a common lysosomal sorting motif (Kienzle and von Blume 2014)(**Fig. 2A**). To understand the impact of the described motifs on GPNMB lysosomal sorting, individual amino acids were mutated, and GPNMB processing, trafficking and secretion was assessed in HEK293 cells. We initially aimed to express the GPNMB constructs in RAW264.7 macrophages, however, the transfection efficiency was too low to generate meaningful results.

**Fig. 2:**
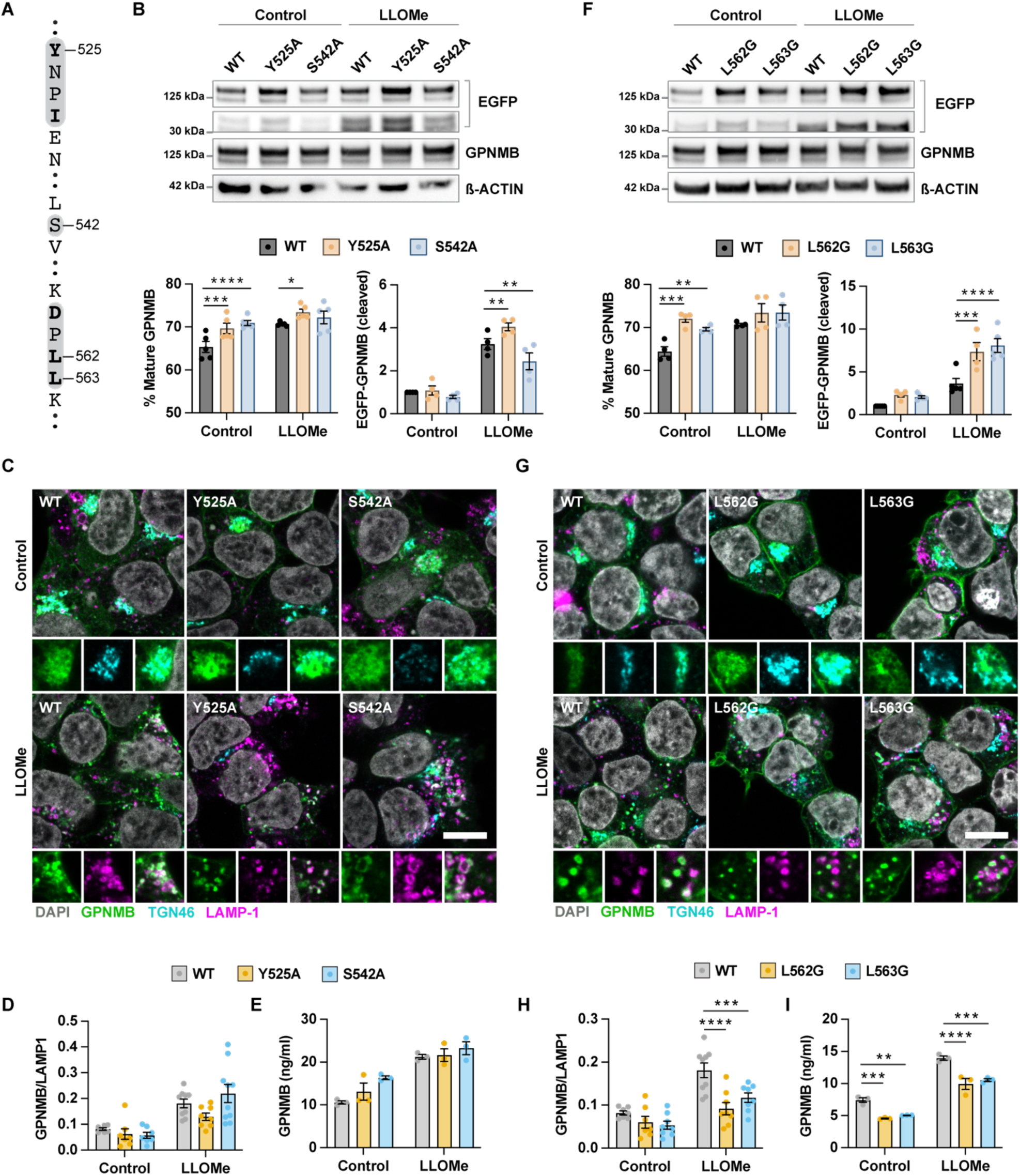
GPNMB lysosomal recruitment is a pre-requisite for secretion. **(A)** Overview of potential lysosomal recruitment motifs in the C-terminal tail of GPNMB including a HemITAM (Y525), a phosphorylated Serine (S542) and a Di-Leucine motif (L562 and L563). **(B-E)** HEK293 cells were transfected with GPNMB-EGFP WT or containing a Y525A or S542A point mutation. **(B)** GPNMB processing was assessed by Western blotting for the C-terminal EGFP-tag and with an N-terminal targeting anti-GPNMB antibody. WBs were quantified using ImageJ. Graphs show mean +/- SEM, n=4 **(C)** GPNMB localisation was imaged by confocal microscopy. Scale bar = 10 μm. **(D)** GPNMB-LAMP-1 co-localisation was quantified using ImageJ. Graphs show mean +/- SEM of n=3 with 2-4 technical replicates each. **(E)** GPNMB secretion was measured by ELISA. Graphs show mean +/- SEM, n=3. **(F-I)** HEK293 cells were transfected with GPNMB-EGFP WT or containing a L562G or L563G point mutation. **(F)** GPNMB processing was assessed by Western blotting for the C-terminal EGFP- tag and with an N-terminal targeting anti-GPNMB antibody. WBs were quantified using ImageJ. Graphs show mean +/- SEM, n=4. **(G)** GPNMB localisation was imaged by confocal microscopy. Scale bar = 10 μm. **(H)** GPNMB-LAMP-1 co-localisation was quantified using ImageJ. Graphs show mean +/- SEM of n=3 with 2-4 technical replicates each. **(I)** GPNMB secretion was measured by ELISA. Graphs show mean +/- SEM, n=3.

Mutating S542 to A or mutating the proposed HemITAM affected the ratio of mature to precursor protein and altered the amount of the N-terminal cleavage product (**Fig. 2B**). However, these mutations had no obvious effect on GPNMB localisation and secretion before or after LLOMe stimulation (**Fig. 2C-E**).

Similar to the HemITAM and S542A mutants, the L562G and L563G mutants showed a higher ratio of mature to precursor form, and markedly increased the presence of the C-terminal cleavage product at steady-state and after LLOMe stimulation (**Fig. 2F**). Moreover, the C-terminal di-leucine motif was required for GPNMB lysosomal localisation. Both the L562G and L563G mutants showed reduced localisation to LAMP1 positive vesicles and instead mis-localised to the plasma membrane in response to LLOMe treatment (**Fig. 2G-H**). In addition, the secretion of the L562G and L563G mutants was reduced, indicating that GPNMB lysosomal recruitment is a prerequisite for GPNMB secretion (**Fig. 2I**).

### GPNMB is secreted via lysosomal exocytosis

These data point to the GPNMB ectodomain being secreted via the lysosome. Lysosomes secret content by fusing with the plasma membrane which results in the translocation of lysosomal membrane protein to the plasma membrane (Néel et al. 2024). Therefore, measuring the abundance of LAMP-1 on the plasma membrane by flow cytometry can serve as a proxy for lysosomal exocytosis. As predicted, LLOMe, Bafilomycin A1 and Nigericin treatment resulted in lysosomal exocytosis whereas MG132 and LPS did not result in lysosomal exocytosis (**Fig. 3A**). As lysosomal content can be released directly or in vesicles, we inhibited intraluminal vesicle formation using GW4869. GW4869 did not affect GPNMB secretion indicating that GPNMB is not secreted in extracellular vesicles (**Fig. 3B**). To determine if lysosomal exocytosis is required for GPNMB secretion, we modulated the activity of the lysosomal Ca2+ efflux channel TRPML1 which acts as a key mediator of lysosomal exocytosis (Tsunemi et al. 2019; Ballabio and Bonifacino 2020). The TRPML1 channel agonist ML-SA1 increased lysosomal exocytosis (**Fig. 3C**), and increased GPNMB secretion (**Fig. 3D**). Vice versa, the TRPML1 channel antagonist ML-SI3 decreased lysosomal exocytosis (**Fig. 3E**), and strongly decreased GPNMB secretion (**Fig. 3F**). We confirmed that the modulation of lysosomal calcium flux did not directly impact the recruitment of GPNMB to lysosomes (**Fig. 3G**) leading to the conclusion that GPNMB is secreted via lysosomal exocytosis.

**Fig. 3:**
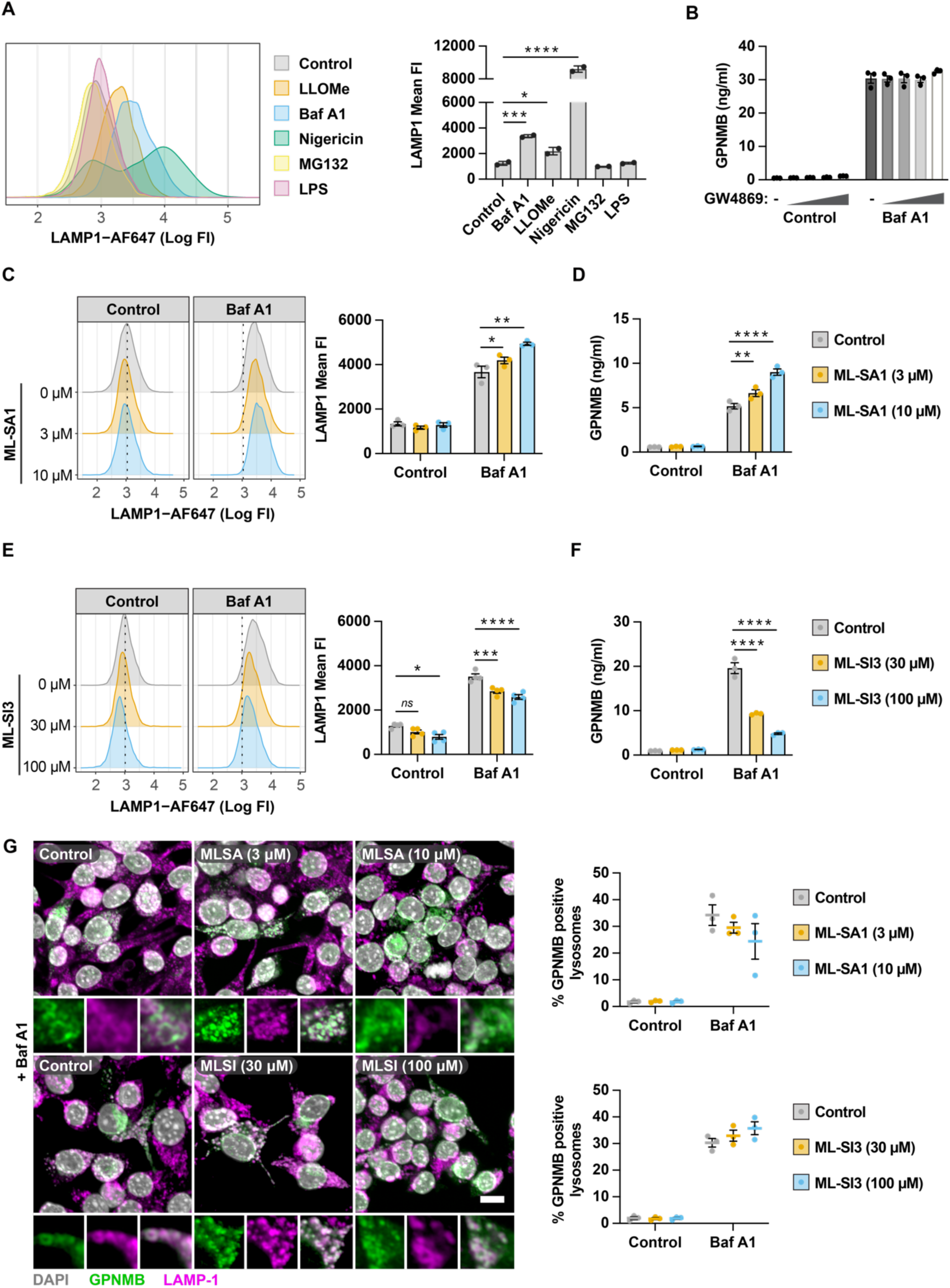
GPNMB is secreted via lysosomal exocytosis. **(A)** Lysosomal exocytosis was estimated by measuring LAMP-1 on the cell surface by flow cytometry. RAW264.7 macrophages were either left untreated or treated with LLOMe (500 μM), Bafilomycin A1 (100 nM), Nigericin (5 μM), MG132 (5 μM) or LPS (10 ng/ml) for 4hrs. Graphs show mean +/- SEM, n=2. **(B)** RAW264.7 macrophages were treated with increasing concentrations of GW4869 (0.3 – 10 μM) and stimulated with Bafilomycin A1 (10 nM) overnight. GPNMB secretion was measured by ELISA. Mean +/- SEM, n=3. **(C)** RAW264.7 macrophages were treated with ML-SA1 and co-stimulated with Bafilomycin A1 (100 nM) for 4 hrs. Lysosomal exocytosis was measured by quantifying the cell surface abundance of LAMP-1 by FACS. Mean +/- SEM, n=3. **(D)** RAW264.7 macrophages were treated with ML-SA1 and co-stimulated with Bafilomycin A1 (10 nM) overnight. GPNMB secretion was measured by ELISA. Mean +/- SEM, n=3. **(E)** RAW264.7 macrophages were treated with ML-SI3 and stimulated with Bafilomycin A1 (100 nM) for 4 hrs. Lysosomal exocytosis was measured by quantifying the cell surface abundance of LAMP-1 by FACS. Mean +/- SEM, n=3. **(F)** RAW264.7 macrophages were treated with ML-SI3 and stimulated with Bafilomycin A1 (10 nM) overnight. GPNMB secretion was measured by ELISA. Mean +/- SEM, n=3. **(G)** RAW264.7 macrophages were treated with ML-SI3 or MLSI and co-stimulated with Bafilomycin A1 (100 nM) for 4 hrs. GPNMB localisation to LAMP-1 positive lysosomes was assessed by high-content imaging. Mean +/- SEM shown, n=3. Scale bar = 10 μm.

### ADAM10 contributes to GPNMB secretion

The secreted N-terminal fragment of GPNMB is thought to be generated by cleavage by α-secretases such as ADAM10 (Furochi et al. 2007). To test the requirement for ADAM10 or ADAM17 for Bafilomycin A1 or LLOMe-stimulated GPNMB secretion, we inhibited ADAM10 and ADAM17 using GW 280264X (Hundhausen et al. 2003). In RAW264.7 macrophages, ADAM10/17 inhibition reduced GPNMB secretion in a dose-dependent manner (**Fig. 4A**). Further, the reduction in mature, fully glycosylated GPNMB due to increased secretion following BafA1 treatment was reversed by ADAM10/17 inhibition (**Fig. 4 B-C**). To further evaluate processing of GPMNB, we sought to detect the C-terminal intracellular fragment in HEK293 cells. In agreement with the findings in RAW264.7 macrophages, ADAM10/17 inhibition reduced the secretion of overexpressed GPNMB in HEK293 cells (**Fig. 4D**).

**Fig. 4:**
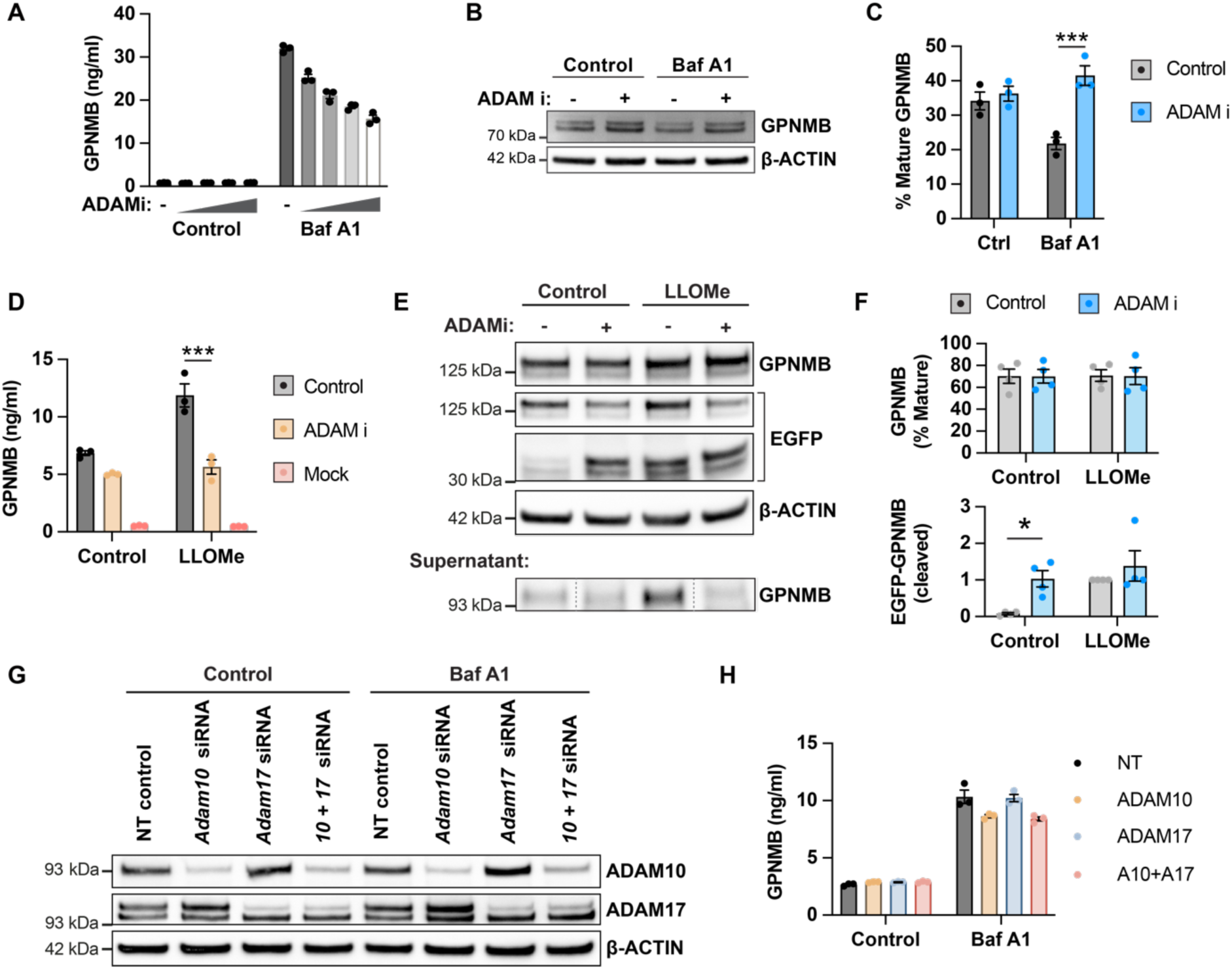
Cleavage by ADAM10 contributes to GPNMB secretion. **(A)** RAW264.7 macrophages were treated with increasing concentrations of the ADAM10/17 inhibitor G280264x (0.3 – 10 μM) and GPNMB secretion in response to Bafilomycin A1 (10 nM, overnight) was measured by ELISA. Mean +/- SEM, n=3. **(B)** RAW264.7 macrophages were pre-treated with cycloheximide (5 μg/ml) and the ADAM10/17 inhibitor G280264x (3 μM) where indicated, followed by stimulation with Bafilomycin A1 (100 nM) for 4 hrs. GPNMB protein levels were measured by Western Blot. **(C)** Quantification of the mature form of GPNMB from (B). Mean +/- SEM, n=3. **(D)** Human GPNMB transfected HEK293 cells were pre-treated with the ADAM10/17 inhibitor G280264x (3μM) and GPNMB secretion in response to LLOMe (1 mM, overnight) was measured by ELISA. Mean +/- SEM, n=3. Mock indicates the non-transfected control. **(E)** Human GPNMB transfected HEK293 cells were pre-treated with the ADAM10/17 inhibitor G280264x (3 μM) and stimulated with LLOMe (1 mM) for 1 hr. GPNMB protein levels in the cell lysate and supernatant were measured by Western Blot. **(F)** Quantification of the mature form of GPNMB and the EGFP-tagged C-terminal tail from B. Mean +/- SEM, n=4.**(G)** RAW264.7 macrophages were treated with 100 nM of siRNAs targeting ADAM10 or ADAM17 and stimulated with Bafilomycin A1 (10 nM) overnight. Knock-down efficiency was assessed by Western Blot. **(H)** GPNMB secretion was measured by ELISA. Mean +/- SEM, n=3.

Lysosomal damage induced by treatment with LLOMe increased the amount of the cleaved C-terminal fragment detected with an anti-EGFP antibody, indicating either increased cleavage or impaired degradation. Surprisingly, ADAM10/17 inhibition increased the amount of the C-terminal fragment at steady-state but did not significantly alter the amount after LLOMe stimulation - supporting a complex interplay between cleavage and degradation (**Fig. 4E-F**). However, in the HEK293 overexpression system, we were able to detect secreted GPNMB in the supernatant by Western Blotting confirming that LLOMe treatment increases secretion which is inhibited by the ADAM10/17 inhibitor (**Fig. 4E**). In line with reduced cleavage and secretion, ADAM10/17 inhibition resulted in increased presence of GPNMB on lysosomes and the cell surface (**Fig. S1**). To differentiate between cleavage by ADAM10 and ADAM17, we knocked-down ADAM10 or ADAM17 in RAW264.7 cells and assessed GPNMB secretion. Despite similar levels of knock-down (**Fig. 4G**), only ADAM10 knock-down reduced GPNMB secretion (**Fig. 4H**). However, we noted that despite a high degree of knock-down, the effect on GPNMB secretion was minimal, indicating that either minimal levels of ADAM10 activity are sufficient for cleavage, or that alternative ADAM secretases or proteases can cleave GPNMB. We concluded that ADAM10 contributes to GPNMB lysosomal secretion, but might not be the sole protease responsible for cleavage.

### The LRRK2 G2019S pathogenic variant increases GPNMB secretion

GPNMB has been proposed as a biomarker for PD, but increased GPNMB serum levels have also been reported in Gaucher’s Disease patients who carry homozygous *GBA1* mutations (Murugesan et al. 2018) and pharmacological inhibition of glucocerebrosidase increased GPNMB protein levels in the brain of WT mice (Moloney et al. 2018). As heterozygous *GBA1* mutations are a prominent risk factor for developing PD, we aimed to address the question if defined monogenic forms of PD present with altered GPNMB processing in the central nervous system of PD patients. We analysed the Parkinson’s Progression Markers Initiative dataset that includes GPNMB CSF levels of idiopathic PD patients, PD patients carrying *GBA1* mutations and PD patients carrying the *LRRK2 G2019S* mutation. GPNMB levels in idiopathic PD patients were slightly elevated when compared to age-matched, healthy controls as previously reported and *GBA1* mutation carriers showed elevated GPNMB levels to a similar degree as iPD patients (**Fig. 5 A**). To our surprise, *LRRK2* mutation carriers showed highly elevated GPNMB CSF levels (**Fig. 5 A**) implying an impact of LRRK2 function on GPNMB biology. The pattern of GPNMB CSF levels paralleled that observed with urinary di-22:6-BMP levels in the same patient cohort (Merchant et al. 2023) (**Suppl. Fig. 1A**). Although LRRK2 activity seems to be driving both the presence of the lysosomal lipid di-22:6-BMP in the urine and the secretion of GPNMB into the CSF, we only observed a poor correlation of GPNMB CSF levels and di-22:6-BMP levels in the urine (**Suppl. Fig. 1B-C**) indicating different mechanisms of release.

**Fig. 5:**
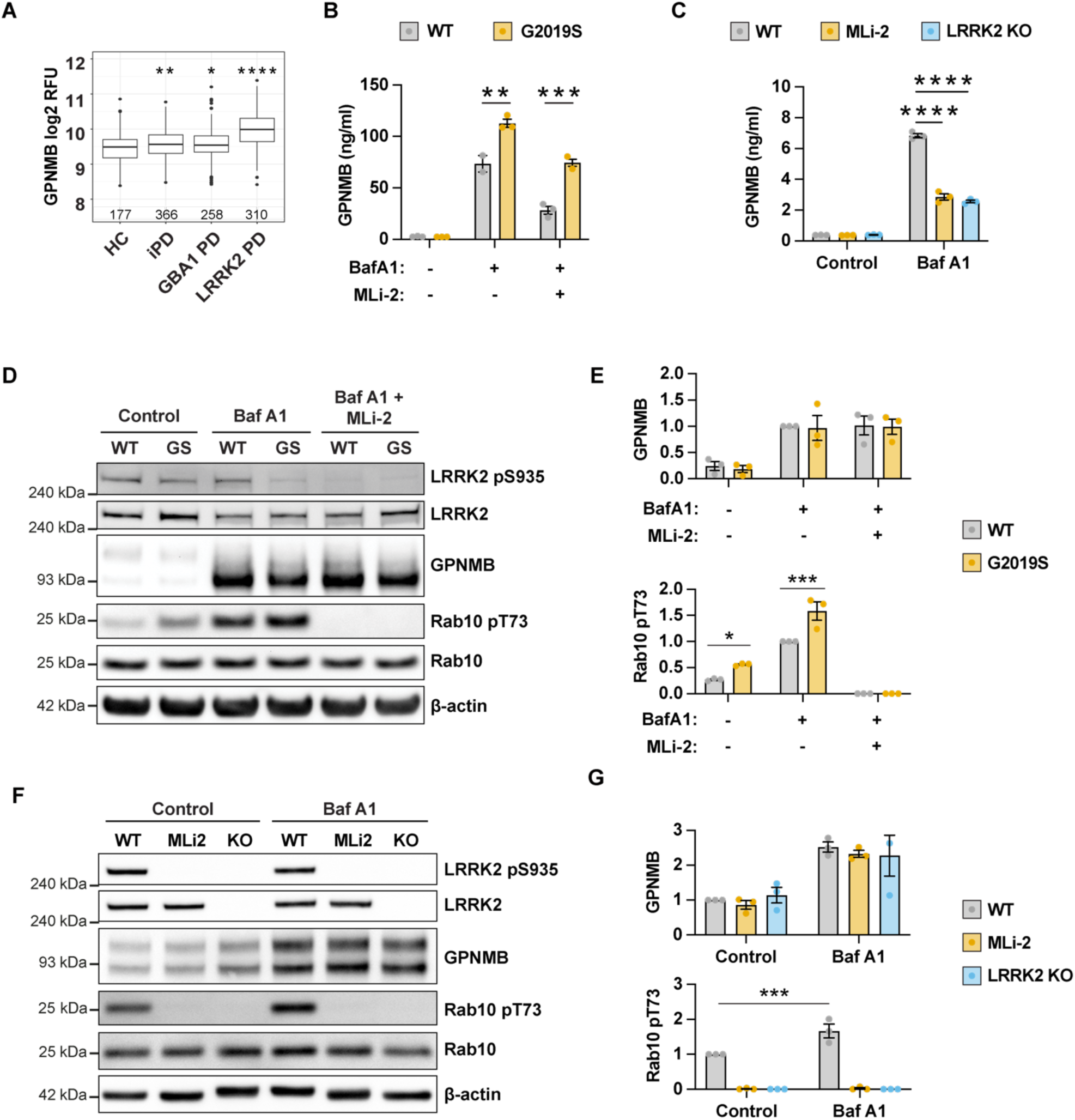
The LRRK2 G2019S mutation increases GPNMB secretion. **(A)** SomaScan data of GPNMB levels in the CSF of healthy controls (HC), idiopathic PD patients (iPD), GBA1 PD patients and LRRK2 G2019S PD patients indicates higher GPNMB levels in LRRK2 mutation carriers. **(B)** WT and LRRK2 G2019S BMDMs were pre-treated with MLi-2 (100 nM) and stimulated with Bafilomycin A1 (10 nM) overnight. GPNMB levels were measured by ELISA. Mean +/- SEM, n=3. **(C)** WT and LRRK2 KO RAW264.7 macrophages were pre-treated with MLi-2 (100 nM) and stimulated with Bafilomycin A1 (10 nM) overnight. GPNMB levels were measured by ELISA. Mean +/- SEM, n=3. **(D)** WT and LRRK2 G2019S BMDMs were pre-treated with MLi-2 (100 nM) and stimulated with Bafilomycin A1 (10 nM) overnight. GPNMB levels and LRRK2 activation was assessed by Western Blotting. **(E)** Quantification of total GPNMB protein levels (premature + mature form) and Rab10 T73 phosphorylation from (D). Graph shows mean +/-SEM, n=3. **(F)** WT and LRRK2 KO RAW264.7 macrophages were pre-treated with MLi-2 (100 nM) and stimulated with Bafilomycin A1 (10 nM) overnight. GPNMB levels and LRRK2 activation was assessed by Western Blotting. **(G)** Quantification of total GPNMB protein levels (premature + mature form) and Rab10 T73 phosphorylation from F). Graph shows mean +/-SEM, n=3.

To verify that LRRK2 activity modifies GPNMB secretion, we stimulated bone marrow-derived macrophages from WT and G2019S LRRK2 knock-in mice with Bafilomycin A1 in the absence or presence of the LRRK2 kinase inhibitor MLi-2 (Fell et al. 2015). LRRK2 kinase gain-of-function resulted in increased GPNMB secretion as was seen in the CSF of LRRK2 G2019S mutation carriers. Vice versa, LRRK2 kinase inhibition strongly inhibited GPNMB secretion, both in WT and G2019S macrophages (**Fig. 5B**). Consistent with kinase inhibition, we also observed reduced GPNMB secretion in LRRK2 KO RAW264.7 macrophages (**Fig. 5C**). LRRK2 has been implicated in a multitude of cellular pathways, including in the regulation of TFEB-regulated transcription (Yadavalli and Ferguson 2023). LRRK2 kinase inhibition or the LRRK2 G2019S mutation had no impact on total GPNMB protein levels (**Fig. 5D-E**) neither had LRRK2 KO in RAW264.7 cells (**Fig. 5F-G**). Western Blotting for the phosphorylation of the LRRK2 substrate Rab10 confirmed LRRK2 kinase inhibition, overactivation or loss-of-function (**Fig. 5D-G**) indicating that LRRK2-mediated alterations in TFEB activity are unlikely to underly alterations in GPNMB secretion.

### LRRK2 modulates GPNMB lysosomal recruitment and lysosomal exocytosis

To gain an insight into how LRRK2 modulates GPNMB secretion, we first confirmed that GPNMB lysosomal recruitment was maintained in the absence of LRRK2 kinase activity. MLi-2 treatment had a minimal impact on GPNMB recruitment in RAW264.7 macrophages (**Fig. 6A-B**) and in BMDMs (**Fig. 6C-D**). However, the LRRK2 KO showed a strong reduction in GPNMB recruitment and a strong reduction in the overall number of cells that showed GPNMB lysosomal recruitment (**Fig. 6A-B**) indicating that a complete loss of LRRK2 might alter the lysosomal stress response. As LRRK2 has been implicated in lysosomal secretion (Eguchi et al. 2018, 2024), we assessed lysosomal secretion by measuring the presence of LAMP-1 on the plasma membrane. LRRK2 KO and LRRK2 kinase inhibition strongly reduced lysosomal exocytosis in RAW264.7 cells (**Fig. 6E**). Vice versa, lysosomal exocytosis was increased in G2019S macrophages, which could be reversed by LRRK2 kinase inhibition (**Fig. 6F**). These results indicate that LRRK2 modulates GPNMB secretion predominantly by modulating lysosomal exocytosis. In summary, we propose a mechanism by which GPNMB gets recruited to lysosomes in response to lysosomal dysfunction. Dysfunctional lysosomes are exocytosed, resulting in the secretion of the GPNMB extracellular domain. Lysosomal exocytosis in enhanced in the presence of LRRK2 gain-of-function mutations (**Fig. 7**).

**Fig. 6:**
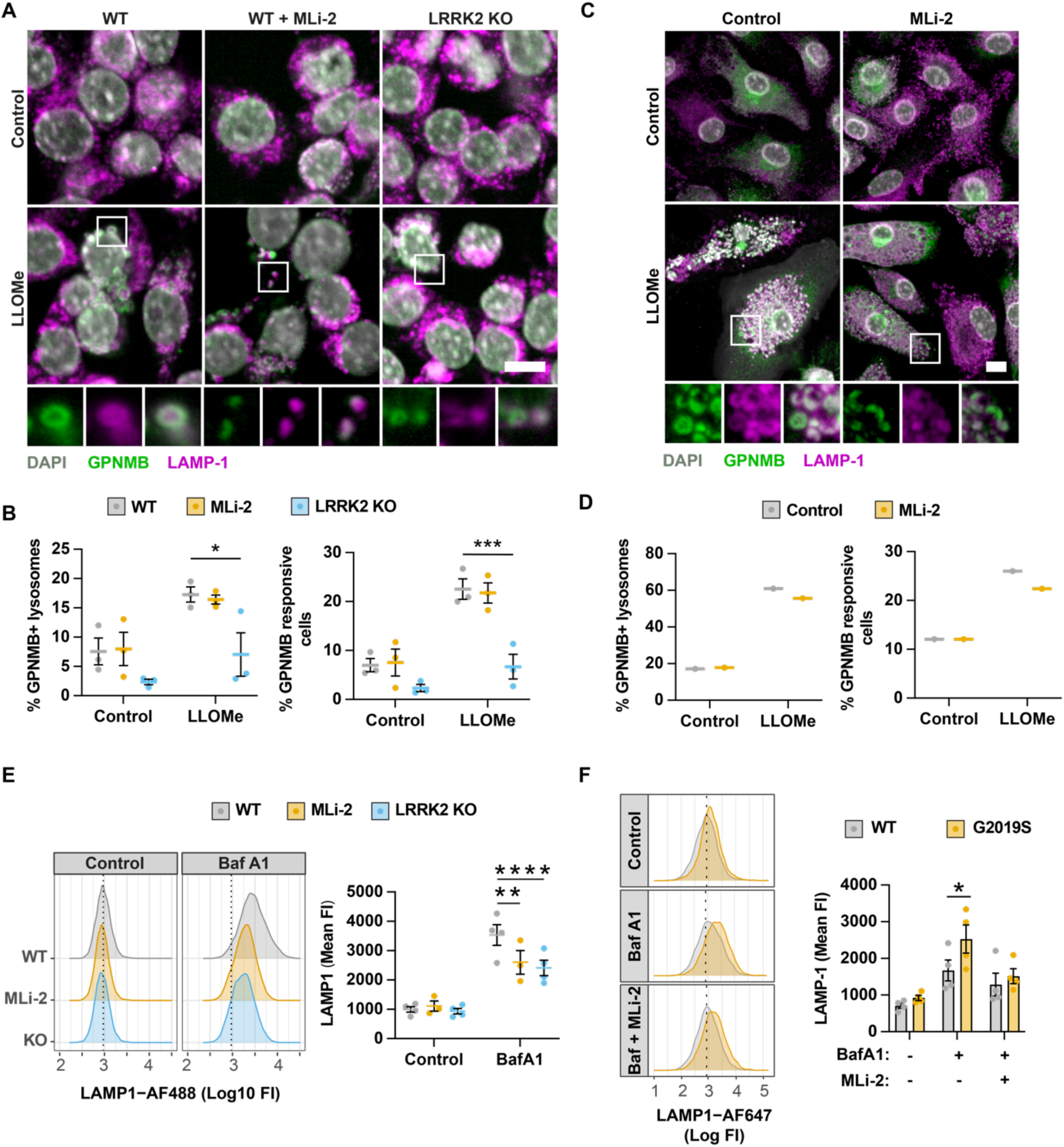
LRRK2 modulates GPNMB lysosomal recruitment and lysosomal exocytosis. **(A)** WT and LRRK2 KO RAW264.7 macrophages were pre-treated with MLi-2 (100 nM) and stimulated with LLOMe (1 mM) for 1hr. GPNMB-LAMP-1 co-localisation was assessed by high-content imaging. Scale bar = 10 μm. **(B)** Quantification of % GPNMB positive lysosomes and % GPNMB responsive cells from (A). Mean +/- SEM, n=3. **(C)** WT BMDMs were pre-treated with MLi-2 (100 nM) and stimulated with LLOMe (1 mM) for 1hr. GPNMB-LAMP-1 co-localisation was assessed by high-content imaging. Scale bar = 10 μm. **(D)** Quantification of % GPNMB positive lysosomes and % GPNMB responsive cells from (C). n = 1. **(E)** WT and LRRK2 KO RAW264.7 macrophages were pre-treated with MLi-2 (100 nM) and stimulated with Bafilomycin A1 (100 nM) for 4hrs. Lysosomal exocytosis was estimated by measuring LAMP-1 on the cell surface by flow cytometry. Graphs show mean +/- SEM, n=3-4. **(F)** WT and LRRK2 G2019S BMDMs were pre-treated with MLi-2 (100 nM) and stimulated with Bafilomycin A1 (100 nM) for 6 hrs. Lysosomal exocytosis was estimated by measuring LAMP-1 on the cell surface by flow cytometry. Graphs show mean +/- SEM, n=4.

**Fig. 7:**
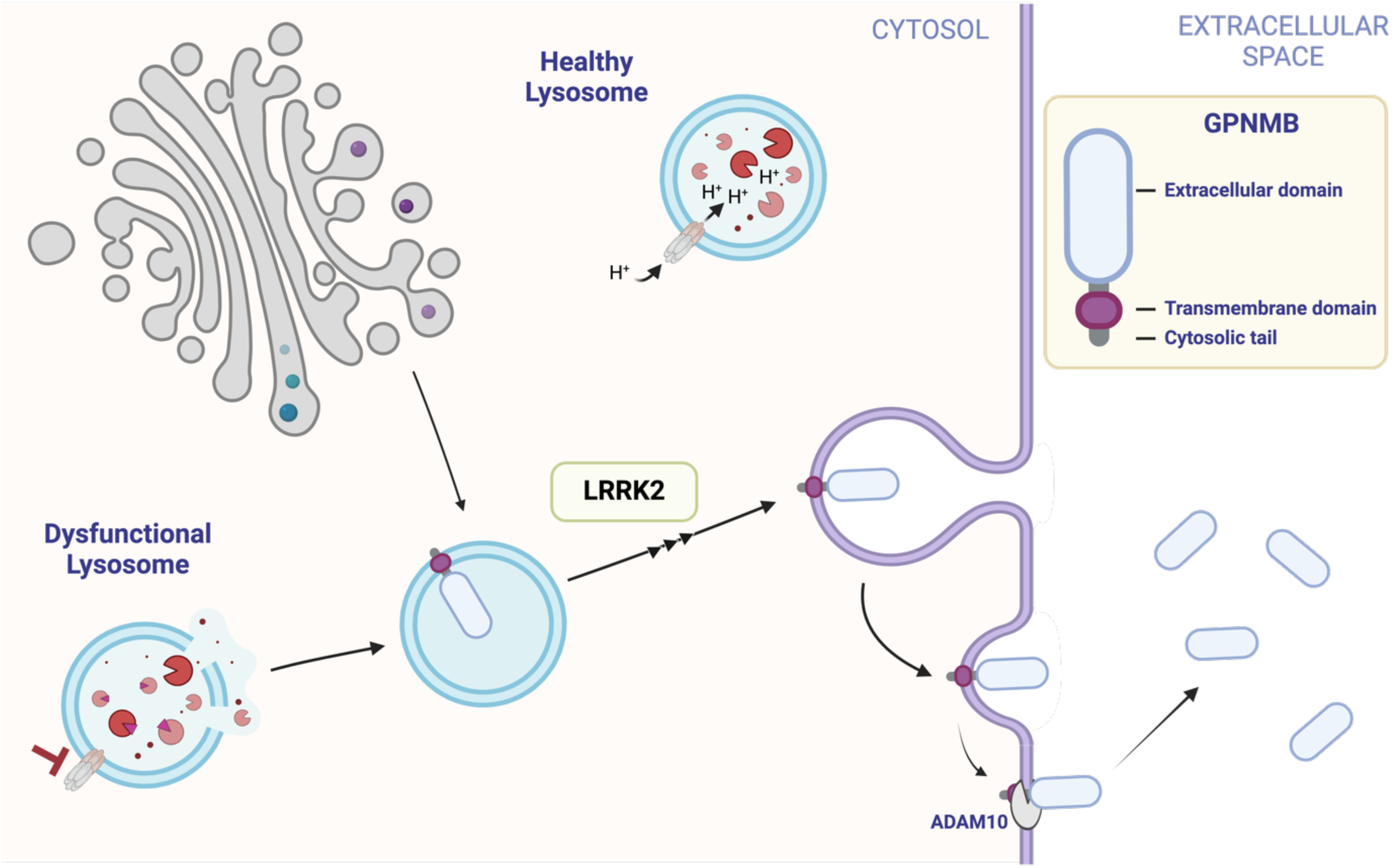
Lysosomal dysfunction results in GPNMB secretion via lysosomal exocytosis. Schematic of the proposed model of GPNMB secretion. GPNMB is not present on healthy lysosomes, but lysosomal dysfunction recruits GPNMB to the lysosomal membrane form the trans-Golgi. The dysfunctional lysosomes get exocytosed, resulting in the presence of GPNMB on the plasma membrane where it gets cleaved for secretion by secretases, including ADAM10.

## Discussion

Due to the association of elevated secreted GPNMB levels to PD risk, GPNMB has emerged as a candidate biomarker or even therapeutic target in Parkinson’s Disease. This study identifies lysosomal dysfunction as a trigger of GPNMB secretion and highlights LRRK2 as a modulator of GPNMB secretion.

GPNMB is functionally associated with a so-called disease-associated microglia phenotype which was initially described in mouse models of Alzheimer’s Disease but similar transcriptional signatures have been validated in human tissue across a multitude of neurodegenerative diseases including Parkinson’s Disease (Keren-Shaul et al. 2017; Paolicelli et al. 2022). Although Smajic *et al*. (Smajić et al. 2022) described GPNMB as a marker for pro-inflammatory microglia in idiopathic Parkinson’s Disease, we also noted that GPNMB expressing microglia display a transcriptional signature of lipid dyshomeostasis and a high degree of expression of lysosome-related transcripts, including the MiT-family transcription factor MiTF, suggesting an ongoing high demand on lysosomal function in these microglia.

Our data indicates that as well as being responsive to lysosomal stress on a transcriptional level, GPNMB secretion is triggered by lysosomal dysfunction. Lysosomal dysfunction results in recruitment of GPNMB to the lysosomal membrane and secretion via lysosomal exocytosis. Although predominantly known for their degradative ability and as a signaling hub (Ballabio and Bonifacino 2020), lysosomes have gained recognition as a secretory compartment. Secreted lysosomes tend to be characterised by a less acidic luminal pH and impaired degrative ability (Néel et al. 2024) implying that lysosomal dysfunction precedes secretion. Coupled with the finding that GPNMB is only present on lysosomes under stress conditions, it is intriguing to speculate that GPNMB represents a “danger” signal, signaling lysosomal stress in macrophages to the surrounding cells.

This study uses pharmacological means to induce lysosomal stress. However, α-synuclein aggregates and tau fibrils have been shown to deacidify the lysosomal lumen and cause lysosomal damage akin to the perturbation used in this study (Rose et al. 2024; Kakuda et al. 2024; Bussi et al. 2018). Further to this, α-synuclein has been shown to be released via lysosomal exocytosis (Tsunemi et al. 2019; Bae et al. 2022) making it likely that microglia experience similar stressors in Parkinson’s Disease.

The signaling events that follow GPNMB secretion remain largely unknown. Secreted GPNMB has been attributed anti-inflammatory but also senescence-inducing properties and is thought to signal via CD44 (Nickl, Qadri, and Bader 2021; Neal et al. 2018; Suda et al. 2021). In the context of Parkinson’s Disease, GPNMB has been highlighted as a receptor for α-synuclein uptake (Diaz-Ortiz et al. 2022). Whether secreted GPNMB therefore can act as an α-synuclein-binding partner requires further investigation. Our findings also highlight an interplay between GPNMB secretion and the Parkinson’s risk gene *LRRK2*. The LRRK2 G2019S gain-of-function mutations increased GPNMB secretion whereas inhibition or loss of LRRK2 kinase function decreased GPNMB secretion substantially. Of note, high GPNMB levels have also been observed in the CSF of carriers of an LRRK2 risk variant that increases PD risk independent of the LRRK2 G2019S mutation (Phillips et al. 2023; Lake et al. 2022) further strengthening the connection between LRRK2 and GPNMB release.

On a functional level, LRRK2 has been shown to mediate lysosomal secretion in response to luminal alkalinization (Eguchi et al. 2018) and is recruited to lysosomes in response to lysosomal stress via unconventional autophagy (Herbst et al. 2020; Eguchi et al. 2024; Bentley-DeSousa and Ferguson 2023; Bonet-Ponce et al. 2020a). LRRK2 phosphorylates a subset of Rab GTPases (Martin Steger et al. 2016), including Rab10, which is directly involved in lysosomal exocytosis (Encarnação et al. 2016), potentially explaining how LRRK2 modulates GPNMB secretion. However, at this point, we can not exclude that the altered secretion is due to LRRK2 directly impacting lysosomal properties such as acidification or the lysosomal stress response (Härtlova et al. 2018; Herbst et al. 2020) which in turn also could impact the degree of lysosomal exocytosis. As elevated GPNMB CSF levels alone increase PD risk, these findings also raise the question if GPNMB contributes to PD risk in LRRK2 mutation carriers.

Taken together, this study highlights GPNMB as a secreted biomarker for lysosomal stress. Although the increased GPNMB levels in idiopathic PD indicate that lysosomal dysfunction is a common feature in Parkinson’s Disease, we postulate that GPNMB will act as a macrophage lysosomal stress marker in a wide spectrum of diseases. The link to LRRK2 biology is intriguing and further highlights the need to understand the downstream consequences of GPNMB secretion.

## Materials and Methods

### Cell culture

Murine bone marrow-derived macrophages were generated by differentiating bone marrow from WT (C57BL/6NTac, Taconic) and LRRK2 G2019S knock-in mice (C57BL/6-Lrrk2tm4.1Arte, Taconic, both kindly provided by Prof Dario Alessi, University of Dundee) in 50 ng/ml murine GM-CSF (576306, BioLegend) in DMEM/10 % FCS for 7 days. RAW264.7 WT (ATCC Cat# SC-6003, RRID:CVCL_UL71) and LRRK2 KO macrophages (ATCC Cat# SC-4 RRID:CVCL_UL72) were obtained from ATCC and cultured in DMEM/10 % FCS. HEK293T cells were obtained from ATCC (ATCC CRL-3216; RRID:CVCL_0063) and cultured in DMEM/10% FCS. If HEK293T cells were seeded on glass coverslips, the coverslips were pretreated with poly-D-Lysine (Gibco, ThermoFisher Scientific). All cells were cultured at 37 <C, 5% CO2.

### Inhibitors and Cell stimulations

Bafilomycin A1 (SML1661, Sigma-Aldrich) was used at 10 nM for ELISA sample preparation or 100 nM for short-term treatment. LLOMe (H-Leu-Leu-OMe HBr, # 4000725, Bachem Biochemica) was used at 500 μM for ELISA sample preparation, or at 1 mM LLOMe for short-term treatments. GW280264 X (# 7030, TOCRIS) was used at 3 μM, GW4869 (# 6741, TOCRIS) was used at 0.3 -10 μM, ML SA1 (# 4746, TOCRIS) and used at 3 – 10 μM, (1R,2R)-ML-SI3 (# HY-134819A, MedChemExpress) was used at 30 – 100 μM, Nigericin (# tlrl-nig, Invivogen) was used 5 μM, the proteasome inhibitor MG-132 (tlrl-mg132, Invivogen) was used at 5 μM and LPS (# L3129, Sigma) was used at 10 ng/ml. The LRRK2 kinase inhibitor MLi-2 (# 5756, TOCRIS) was used at 100 nM.

### Plasmids and site-directed mutagenesis

The open reading frame of full-length human GPNMB (NM_001005340.2) cloned into pcDNA3.1-C-EGFP was purchased from GenScript. GPNMB mutant constructs were generated by site-directed mutagenesis using the Q5 SDM kit from New England Biolabs (#E0552S, dx.doi.org/<u>10.17504/protocols.io.bddfi23n</u>). All plasmids were verified by sequencing. All DNA constructs were maintained in *Escherichia coli* DH5α (#11583117, Thermo Fisher Scientific) and extracted using a plasmid miniprep kit from Qiagen.

### Antibodies

Antibodies used for Western Blotting and immunofluorescence were goat-anti-human-GPNMB (R&D Systems Cat# AF2550, RRID:AB_416615), goat-anti-mouse-GPNMB (AF2330, R&D Systems), mouse-anti-human-LAMP1 (DSHB Cat# h4a3, RRID:AB_2296838), rabbit-anti-mouse-LAMP1 (Abcam Cat# ab208943, RRID:AB_2923327), rabbit-anti-TGN46 (Proteintech Cat# 13573-1-AP, RRID:AB_10597396), mouse-anti-GFP (Thermo Fisher Scientific Cat# MA5-15256, RRID:AB_10979281)), rabbit-anti-ADAM10 (Abcam Cat# ab124695, RRID:AB_10972023), rabbit-anti-ADAM17 (Proteintech Cat# 29948-1-AP, RRID:AB_2935490), rabbit-anti-LRRK2 (Abcam Cat# ab133474, RRID:AB_2713963), rabbit-anti LRRK2 pS935 (Abcam Cat# ab133450, RRID:AB_2732035), rabbit-anti-Rab10 pT73 (Abcam Cat# ab241060, RRID:AB_2884876), rabbit-anti-Rab10 (Abcam Cat# ab237703, RRID:AB_2884879) and mouse-anti-β-actin (Sigma-Aldrich Cat# A1978, RRID:AB_476692). Rat-anti-mouse-LAMP-1-AF647 (BioLegend Cat# 121610, RRID:AB_571990) was used for FACS.

### Immunofluorescence microscopy

Cells seeded on coverslips were fixed with 4% methanol-free PFA (15710, Electron Microscopy Sciences) in PBS for 15 min at 4°C. The samples were permeabilised and blocked with 0.3% Triton X-100, 5% FCS in PBS for 20 min. Primary antibodies were diluted in PBS containing 5% FCS and incubated for 1 hr at RT. The samples were washed three times in PBS, and incubated with the secondary antibody diluted in 5% FCS in PBS (anti-goat, anti-mouse or anti-rabbit-Alexa fluor 488, Alexa fluor 568 or Alexa fluor 647, Invitrogen) for 45 min at room temperature. After three more washes with PBS, nuclear staining was performed using 300 nM DAPI (Life Technologies, D3571) in PBS for 10 min. One final wash with PBS was performed before mounting the coverslips on glass slides using DAKO mounting medium (DAKO Cytomation, S3023). Images were acquired on a Leica SP8 inverted microscope or an OPERA Phenix high-content imaging system. A detailed protocol can found under http://dx.doi.org/10.17504/protocols.io.4r3l22yz4l1y/v1.

### GPNMB ELISA

Supernatants for ELISA were prepared by incubating cells with stimuli for ∼18 hrs. ELISAs for human GPNMB (DY2550, R&D Systems) and mouse GPNMB (DY2330, R&D Systems) were conducted according to the manufacturer’s instructions. GPNMB release was controlled for total cell numbers per well by staining nuclei directly after sample harvest with Hoechst for 15 min before imaging on a Tecan Spark plate reader. In the case of overexpressed GPNMB, EGFP fluorescence was measured in addition to Hoechst to confirm equal transfection efficiency across constructs.

### Western Blotting

Cells were washed once with PBS and lysed in a 1% Triton-X/Tris-HCl lysis buffer (9803S, Cell Signaling) containing protease and phosphatase inhibitors (#78440, ThermoFisher Scientific). The samples were denatured at 80°C for 8 min in LDS sample buffer and reducing agent (NuPAGE, Life Technologies) and run on a NuPAGE 4–12% Bis-Tris gel (Life Technologies). The gels were transferred onto a PVDF membrane using the TurboBlot transfer system (BioRad). The membranes were blocked in 5% semi-skinned milk in TBS-T (TBS, 0.1% Tween-20) and incubated with primary antibodies in 5% semi-skinned milk in TBS-T at 4°C overnight, followed by incubation with the secondary antibodies in 5% skimmed milk in TBS-T for 1 h at room temperature. Western blots were imaged using the iBright imaging system (Thermo Fisher) and quantified by densitometry using Fiji (RRID:SCR_002285; http://fiji.sc) (Schindelin et al. 2012). A detailed protocol can be found under http://dx.doi.org/10.17504/protocols.io.4r3l22yz4l1y/v1.

### FACS analysis of lysosomal exocytosis

Cells were treated with Bafilomycin A1 at 100 nM for 4 - 6 hrs and harvested in PBS. Fc receptor binding was blocked using a CD16/CD32 targeting antibody (BD Biosciences Cat# 553142, RRID:AB_394657) diluted in 5 % FCS/PBS for 10 min on ice. The LAMP1-AF647 antibody was added at 0.5 μl per sample for 30 min on ice, followed by dead/live stain (Fixable Viability Dye eFluor-450, # 65-0863-18, eBioscience) for 10 min on ice. The samples were fixed in 2 % PFA/PBS and acquired on a BD FACSCanto^TM^ flow cytometer. Data were analysed and plotted using the CytoExploreR package for R (Hammill 2021). A detailed protocol for staining can be found under DOI: dx.doi.org/10.17504/protocols.io.n92ld8oyxv5b/v1.

### PPMI SomaScan data analysis

A CSF Somascan dataset was obtained on 2023-01-27 from the Parkinson’s Progression Markers Initiative (PPMI) database (RRID:SCR_006431, project ID 151). The dataset was filtered to only contain baseline data and remove participants labelled as prodromal. Parkinson’s Disease patients were further sub-categorised into “GBA1 PD” and “LRRK2 PD”. “GBA1 PD” consists of carriers of *GBA1* E326K, N370S, T369M or L444P variants, the IVS2+1G>A splice donor variant and the L29Afs*18 loss-of-function variant, thereby including *GBA1* variants that are characterized either mild or severe in their impact as an underlying cause for Parkinson’s Disease (Parlar et al. 2023). “LRRK2 PD” includes LRRK2 G2019S mutation carriers only. LRRK2 mutation carriers that also carry *GBA1* variants were removed from the dataset. As a result, we analysed 1111 participants (44.5 % female, 55.5 % male) for GPNMB extracellular domain CSF levels (Somascan testname = “ 5080-131_3”). Statistical significance between groups was determined by a Kruskal-Wallis test, followed by paired comparisons against the control group using Wilcoxon test. All data was processed and analysed using R (Version 4.3.1 (2023-06-16)) (R Core Team 2023).

### Statistical analysis

Unless otherwise stated, statistical analysis was carried out using Graph Pad Prism software. Multiple comparisons were calculated by two-way ANOVA, followed by Dunnett’s multiple comparison test to test for differences to control. \**p<0.05*, \*\**p<0.01*, \*\*\**p<0.001*, \*\*\*\**p<0.0001*.

### Data and materials availability

Primary data and key lab materials used and generated in this study listed in a Key Resource Table can be viewed at doi: 10.5281/zenodo.14217424.

## Acknowledgments

This research was funded by Aligning Science Across Parkinson’s (ASAP 000478) through the Michael J. Fox Foundation for Parkinson’s Research (MJFF). For the purpose of open access, the authors have applied a CC BY public copyright license to all Author Accepted Manuscripts arising from this submission. PAL is a Royal Society Industry Research Fellow in partnership with LifeArc technologies (IF\R2\222002). We are grateful for LRRK2 G2019S KI mice from Professor Dario Alessi (University of Dundee, UK). We acknowledge PPMI – a public-private partnership – which is funded by the Michael J. Fox Foundation for Parkinson’s Research and funding partners, including 4D Pharma, Abbvie, AcureX, Allergan, Amathus Therapeutics, Aligning Science Across Parkinson’s, AskBio, Avid Radiopharmaceuticals, BIAL, BioArctic, Biogen, Biohaven, BioLegend, BlueRock Therapeutics, Bristol-Myers Squibb, Calico Labs, Capsida Biotherapeutics, Celgene, Cerevel Therapeutics, Coave Therapeutics, DaCapo Brainscience, Denali, Edmond J. Safra Foundation, Eli Lilly, Gain Therapeutics, GE HealthCare, Genentech, GSK, Golub Capital, Handl Therapeutics, Insitro, Jazz Pharmaceuticals, Johnson & Johnson Innovative Medicine, Lundbeck, Merck, Meso Scale Discovery, Mission Therapeutics, Neurocrine Biosciences, Neuron23, Neuropore, Pfizer, Piramal, Prevail Therapeutics, Roche, Sanofi, Servier, Sun Pharma Advanced Research Company, Takeda, Teva, UCB, Vanqua Bio, Verily, Voyager Therapeutics, the Weston Family Foundation and Yumanity Therapeutics.

## Supplementary Materials

**Fig. S1:**
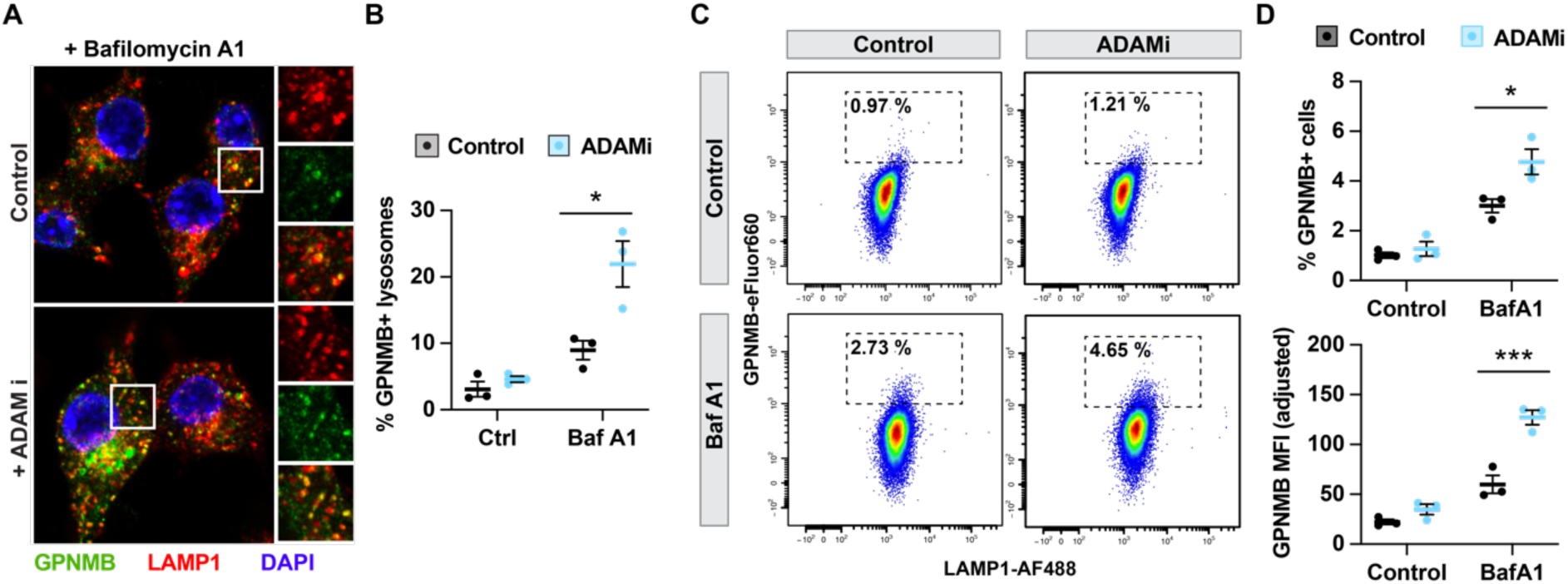
ADAM10/17 inhibition increases GPNMB lysosomal and cell surface localisation. RAW264.7 macrophages were pretreated with 3 μM ADAM10/17 inhibitor, followed by a 4 hr stimulation with 100 nM Bafilomycin A1. **(A)** GPNMB lysosomal co-localisation with the lysosomal marker LAMP-1 was assessed by high-content imaging. **(B)** Quantification of (A) showing mean % of GPNMB+ lysosomes +/- SEM of three independent experiments. **(C)** GPNMB cell surface localisation was assessed by flow cytometry. **(D)** Quantification of (C) showing mean +/- SEM of three independent experiments.

**Fig. S2:**
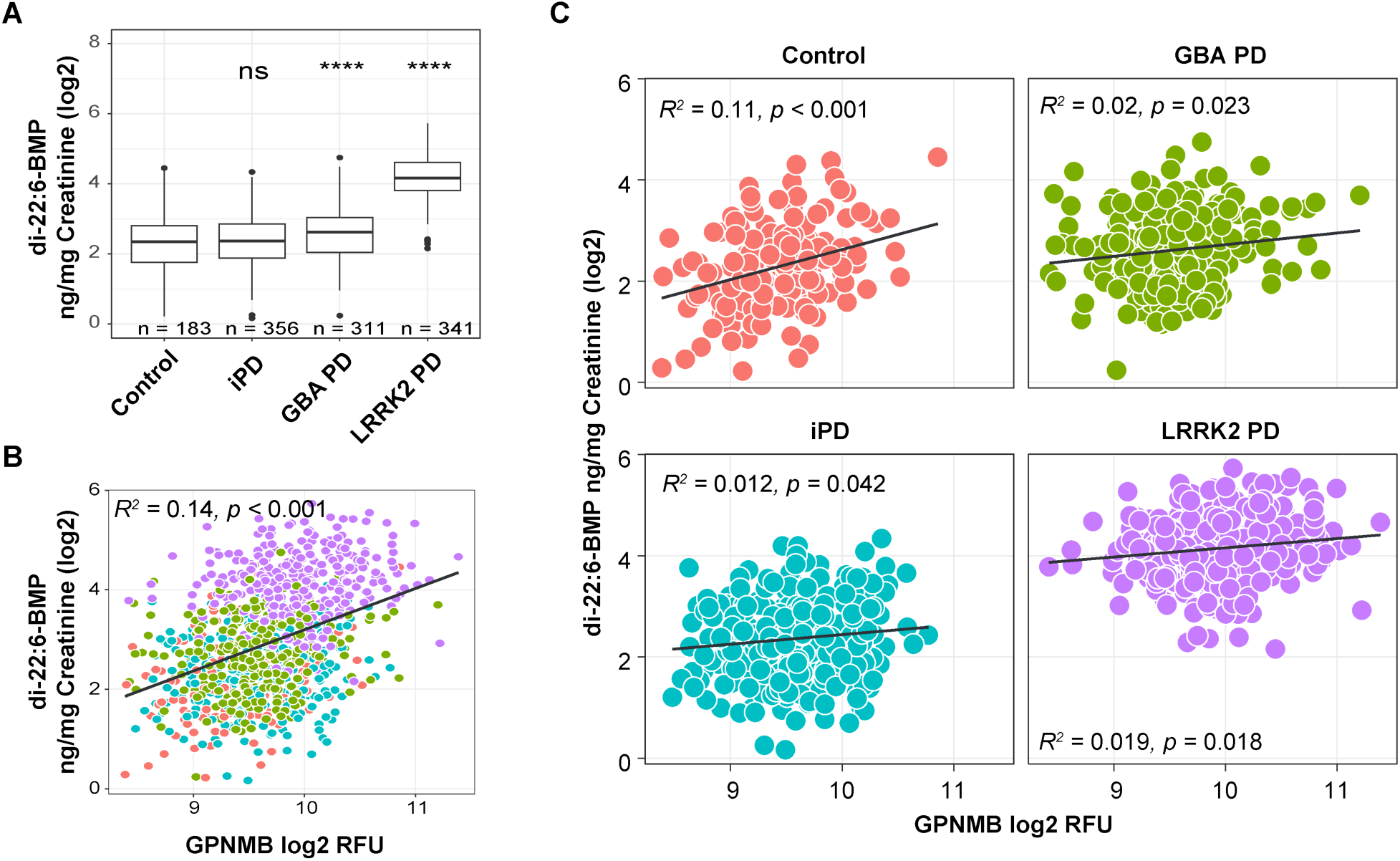
GPNMB CSF and di-22:5-BMP urine levels do not correlate. **(A)** Urine di-22:6-BMP levels in healthy controls (HC), idiopathic PD patients (iPD), GBA1 PD patients and LRRK2 G2019S PD patients indicate higher di-22:5-BMP levels in LRRK2 mutation carriers. **(B)** Correlation of GPNMB CSF levels and di-22:5-BMP urine levels. **(C)** Same data as (B) split by healthy controls, iPD, GBA PD and LRRK2 PD. The colours in (B) and (C) correspond to each other.

